# Phage-display immunoprecipitation sequencing reveals distinct antibody repertoire patterns across a rural-urban gradient

**DOI:** 10.1101/2025.08.26.671278

**Authors:** Ross F. Laidlaw, Gabriel Innocenti, Thomas Vogl, Moustapha Mbow, Maria Yazdanbakhsh, Mikhael D. Manurung

## Abstract

Environmental microbial exposures are hypothesized to drive immune profile differences across rural-urban gradients, yet direct evidence linking exposures to immune phenotypes remains limited. Here, we employed phage-displayed immunoprecipitation sequencing (PhIP-seq) to profile antibody repertoires against 357,192 epitopes from >700 organisms in 62 age-matched healthy adults from rural Senegal (n=29), urban Senegal (n=20), and urban Netherlands (n=13). PhIP-Seq analysis identified two major patterns of antibody repertoire variation: geographically restricted signatures distinguishing Senegal from the Netherlands, and a trajectory associated with rural-urban gradients spanning from rural Senegal to urban Netherlands. Integration of PhIP-seq data with high-dimensional immunological profiling revealed immune signatures correlated with the rural-urban antibody repertoire trajectory. Our findings demonstrate that antibody repertoire profiling captures distinct microbial exposure histories across both geographic regions and rural-urban gradients and correlates with variation in immune signatures. This work establishes PhIP-seq as a powerful tool for linking environmental exposure histories to population-level immune landscapes and provides insights into immune adaptation across diverse environments.

## Introduction

Environmental exposures and lifestyle factors drive variation in immune profiles (1–4) both between geographical regions (5) and across rural-urban gradients within countries (3), influencing disease susceptibility and vaccine responses (6). Understanding these immune variations is critical for developing tailored interventions (7), yet the specific environmental drivers and their mechanisms remain poorly defined.

Emerging evidence reveals multiple contributing factors. Gut microbiome composition varies markedly between rural and urban populations, with rural Tanzanians exhibiting greater microbial diversity and enrichment of *Prevotella* compared to urban populations with higher *Bacteroides* abundance (8, 9) reflecting broader differences between low- and middle-income countries (LMICs) versus high-income countries (HICs) (8). Lifestyle scores also correlate with immune phenotypes, as rural-associated lifestyle scores predict more activated immune profiles than urban-associated scores (10). Additionally, dietary transitions from traditional to Western diets promote inflammatory status (11).

Specific microbial exposures profoundly shape immune profiles through diverse mechanisms. Repeated malaria infections drive distinct transcriptomic profiles in immune cells of Kenyan children (12–14), while *Mycobacterium* exposure—both through BCG vaccination and environmental encounters—remodels innate immune responses (15). Fungal β-glucan exposure enhances monocyte responses to unrelated pathogens (16), and helminth infections selectively expand type 2 and regulatory T and B cell populations (17). These exposure-driven immune remodeling extend beyond LMIC settings: cytomegalovirus-seropositive individuals in HICs exhibit reduced vaccine responses compared to seronegative counterparts, exemplified by diminished Ebola vaccine immunogenicity in exposed versus naïve British adults (10).

Current methodological limitations have precluded direct assessment of microbial exposure differences across rural-urban gradients at the individual level. Phage-displayed immunoprecipitation sequencing (PhIP-seq) now enables scalable, high-resolution profiling of individual antibody repertoires as molecular records of microbial encounters (18–20). This approach has been used to investigate autoimmunity (12–14), microbiome-directed responses (21), and population-level antibody diversity in Northern Netherlands (22), yet its potential for comparing global populations remains unrealized.

Here, we employed PhIP-seq to profile antibody repertoires across individuals residing in rural Senegal, urban Senegal, and urban Netherlands, revealing distinct population-specific antibody epitope patterns. Furthermore, integration with comprehensive multi-omics datasets demonstrate shared variation between antibody repertoires and baseline immune signatures.

This work provides insights into how environmental microbial exposures histories shape immune landscapes across rural-urban gradients in understudied populations.

## Results

### Study population

To investigate antibody repertoire profiles along the rural-urban gradient, we analyzed 62 age-matched healthy adults from rural Senegal (n=29), urban Senegalese (n=20), and urban Netherlands (n=13) (Fig. 1A). Participants showed no significant age differences across groups (median 27 years, range 18-43 years; Kruskal-Wallis *P*=0.55). Sex distribution was balanced within Senegalese cohorts (rural: 48.3% female; urban: 50.0% female) but skewed toward females in the Dutch cohort (76.9% female; Chi-square *P*=0.069). All Senegalese participants had maintained residence in their respective environments for ≥10 years. As described previously (7), rural Senegalese participants exhibited typical rural lifestyle features, including larger household sizes and traditional housing with earth floors and mud walls compared to their urban counterparts.

**Figure 1.**
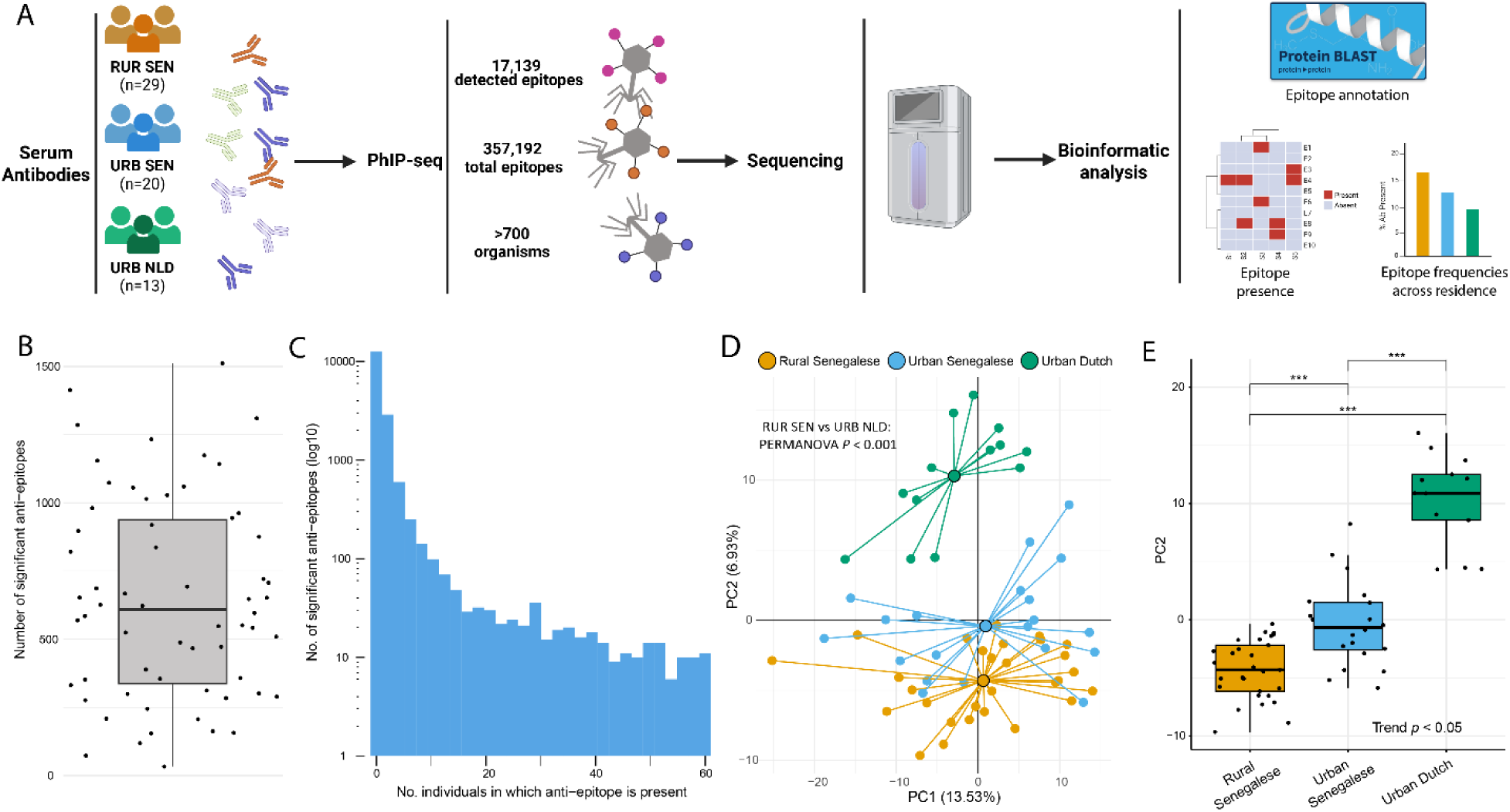
Overview of experiment and PhIP-seq data **A,** Graphical overview of the experiment. Serum from individuals residing in rural and urban Senegal and urban Netherlands was taken. The serum was incubated with a peptide phage library, containing 357,192 epitopes, and immunoprecipitation was performed to capture phages bound by antibodies. The samples were sequenced before bioinformatic analysis to annotate the epitopes (using protein BLAST) and extract patterns of antibody repertoire across the three groups. Created using Biorender.com. **B,** Boxplot showing the number of significant anti-epitopes found per individual. Centre line shows the median and box limits show the 25th and 75th percentiles. The whiskers extend to 1.5 times the interquartile range from the 25th and 75th percentiles. **C,** Histogram of the number of individuals an anti-epitope was detected in. The Y-axis is log10 transform and shows the number of anti-epitopes. **D,** PCA plot generated from the filtered PhIP-seq dataset, where individuals are coloured by residence. The large circles represent the centroid of the different groups in the PC1 and PC2 space. *P* values denote group centroid comparisons with the PERMANOVA test. **E**, Boxplot showing the PC2 values of each individual, grouped by residence. Linear trend test and Tukey post hoc test for pairwise comparisons *P* values are indicated. \*\*\**P*/FDR < 0.001.

### Antibody repertoires vary across the rural-urban gradient

PhIP-seq analysis detected antibodies against 17,139 epitopes from over 700 organisms across all samples (Fig.1A, Supplementary File 1). Individual antibody repertoires contained a median of 609 significant anti-epitope responses (Fig.1B), with substantial heterogeneity in epitope recognition: 12,647 epitopes were detected in at most 1 individual, while 60 epitopes were recognized by over 50 individuals (80%) (Fig. 1C). Total anti-epitope numbers did not differ significantly between groups (Supplementary Fig. 1).

Principal component analysis (PCA) revealed significant differences in overall antibody repertoires between Dutch and rural Senegalese populations (PERMANOVA *P*<0.001). Notably, we observed that rural Senegalese, urban Senegalese, and urban Dutch populations were ordered sequentially along the second principal component (PC2) (Figs. 1D, E), suggesting an underlying rural-urban trajectory of microbial exposure histories. Age and sex showed no significant associations with antibody profiles (Supplementary Figs. 2A, B).

Hierarchical clustering of antibody repertoire profiles revealed several exposure history clusters across the residence groups (Fig. 2A). Clusters 2 and 4 showed geographically restricted signatures, with epitopes enriched mostly in Senegalese or Dutch participants, respectively.

**Figure 2.**
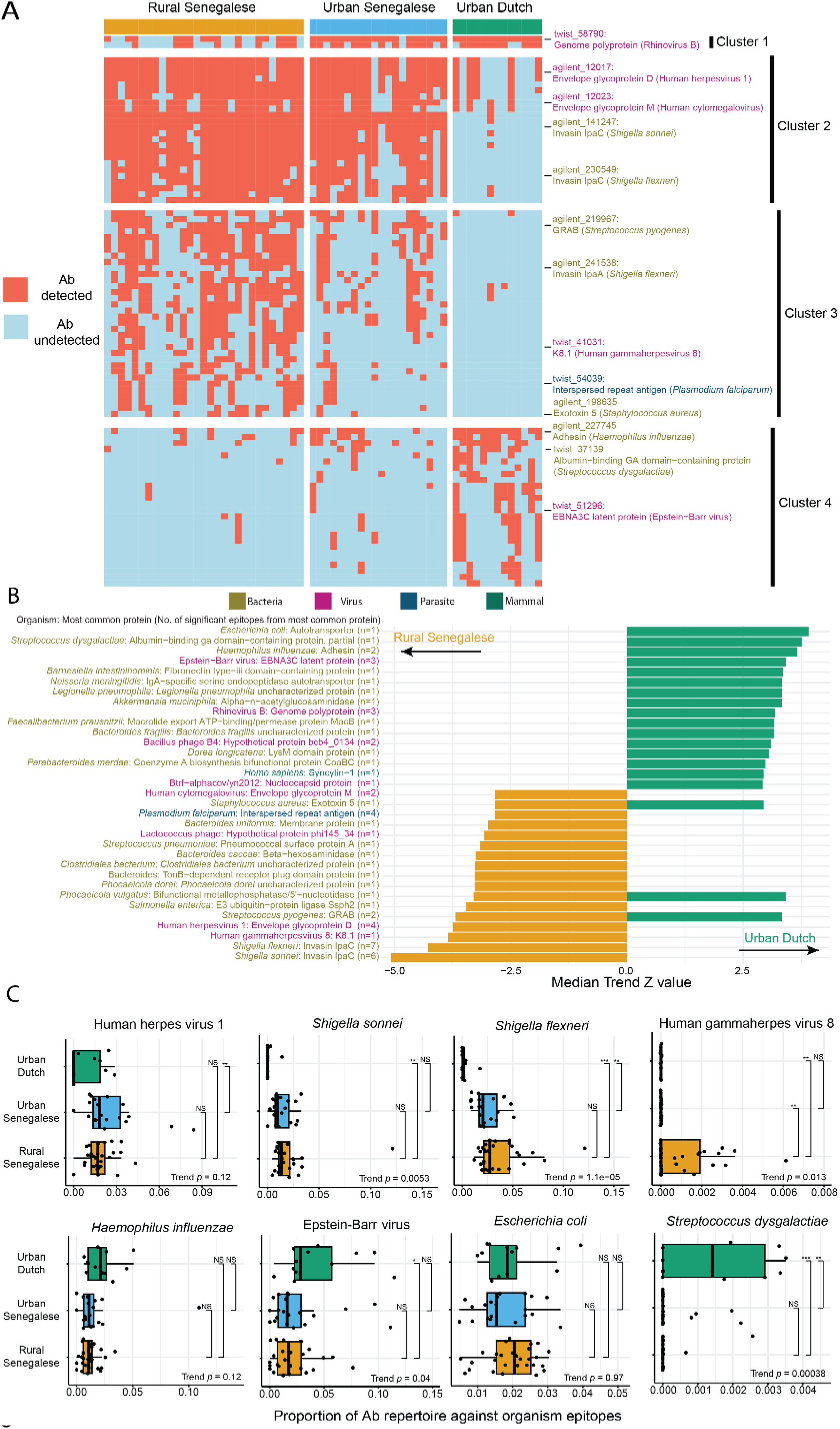
Antibody repertoire analysis across the rural-urban axis. A, Clustered heatmap showing whether antibody (Ab) against a particular epitope was detected (red) or undetected (blue) in an individual. The heatmap is sliced column wise by the residence of the individual, and the rows sliced by k-means clustering on the epitopes. Selected epitopes are annotated on the rows and are coloured according to the type of organism. **B,** Barplot showing the percentage of individuals per residence group who have antibodies against epitopes found to have a significantly linear trend across the rural-urban axis. For each epitope, the protein and organism of origin is detailed. The organism text is coloured by organism type, in the same fashion as panel A. **C,** Barplot showing the percentage of epitopes from a particular organism that were found to have a significant linear trend across the rural-urban axis. The direction and colour of the bars denotes whether the anti-epitopes frequencies trend towards the rural Senegalese or urban Dutch direction. The organism names are coloured by whether they are viral, bacterial, parasitical or mammalian, and the most common protein of the significant anti-epitopes are also denoted. Centre line shows the median and box limits show the 25th and 75th percentiles. The whiskers extend to 1.5 times the interquartile range from the 25th and 75th percentiles. The number of significant anti- epitopes for the most common protein (per organism) is also shown. Linear trend test and Tukey post hoc test for pairwise comparisons P values are indicated. Linear trend p-values adjusted for multiple comparisons using Benjamini-Hochberg. NS *P*/FDR >= 0.05, \**P*/FDR < 0.05, \*\**P*/FDR < 0.01, and \*\*\**P*/FDR < 0.001.

Cluster 3 demonstrated a clear rural-urban gradient, showing highest detection frequencies in rural Senegalese, followed by urban Senegalese, and lowest in urban Dutch subjects.

To quantify these observed clustering patterns, we performed linear trend analysis across the rural-urban gradient. Eighty-six anti-epitopes showed significant rural-urban associations: 28 increased in frequency from rural Senegal to urban Netherlands, while 58 decreased (Fig. 2B, Supplementary Figs. 3-5, Supplementary File 2). These results demonstrate that differences in microbial exposure histories resulted in distinct antibody repertoire signatures spanning both geographic areas and rural-urban gradients.

Pathogen-specific antibody patterns varied across the rural-urban gradient. Rural Senegalese populations showed highest detection for *Shigella spp.*, herpes simplex virus, and cytomegalovirus (CMV) epitopes, while urban Dutch individuals exhibited highest detection for rhinovirus B, Epstein-Barr virus (EBV), and *Haemophilus influenzae* (Fig. 2B). Interestingly, antibodies against epitopes from *Staphylococcus aureus* (*S. aureus*), *Phocaeicola vulgatus* and *Streptococcus pyogenes* proteins showed opposing patterns across the gradient (Fig. 2B), with *S. aureus* protein A anti-epitopes ranking among the most prevalent repertoires (detected in 61/62 individuals; Supplementary File 1).

Since individual epitopes from the same pathogen sometimes showed divergent trends along rural-urban gradients, we assessed pathogen-level enrichment by calculating the proportion of each repertoire targeting organisms with significant linear associations (Supplementary File 2). Analysis of the four organisms with highest median trend Z-values for each direction of association confirmed epitope level findings: *Shigella sonnei, S. flexneri,* and human gammaherpesvirus 8 showed significant rural-trending responses, while EBV and *Streptococcus dysgalactiae* trended toward urban Dutch subjects (Fig. 2C). Geographic restriction in anti-epitope detection was particularly striking—nearly all Senegalese individuals (48/49) recognized *Shigella sonnei* epitopes compared to only three Dutch participants. Human gammaherpesvirus 8 were detected exclusively in rural Senegalese (14/29 individuals) but were completely absent from urban Senegalese and Dutch groups (Fig. 2C).

### PhIP-seq anti-epitope trajectory was correlated with immune-signatures of urbanization

Having established distinct antibody profiles across residence groups, we investigated whether these exposure signatures correlate with immune activation. IgG1 Fc galactosylation, which decreases with immune activation across populations (23), showed a clear rural-urban gradient: lowest in rural Senegalese, intermediate in urban Senegalese, and highest in urban Dutch participants (Fig. 3A). This pattern suggests increasing immune activation from urban to rural settings. Importantly, IgG1 galactosylation levels showed strong negative correlation with the PhIP-seq PC2 scores (Spearman’s ρ=−0.64, *P*<0.0001), suggesting that population-specific microbial exposure histories modulate immune We next integrated PhIP-seq data with comprehensive immune profiling data from the same individuals (Fig. 3B; n = 18 rural Senegalese, 8 urban Senegalese, 4 urban Dutch). Our previously established rural-urban immune trajectory lambda metric (7)—derived from unsupervised integration of multiple immunological datasets—correlated significantly with PhIP-seq PC2 scores (Spearman’s ρ=0.55, *P*=0.0021; Fig. 3C, Supplementary Fig. 6A). This convergence demonstrates that microbial exposure histories captured by antibody repertoires align with immune signatures.

**Figure 3.**
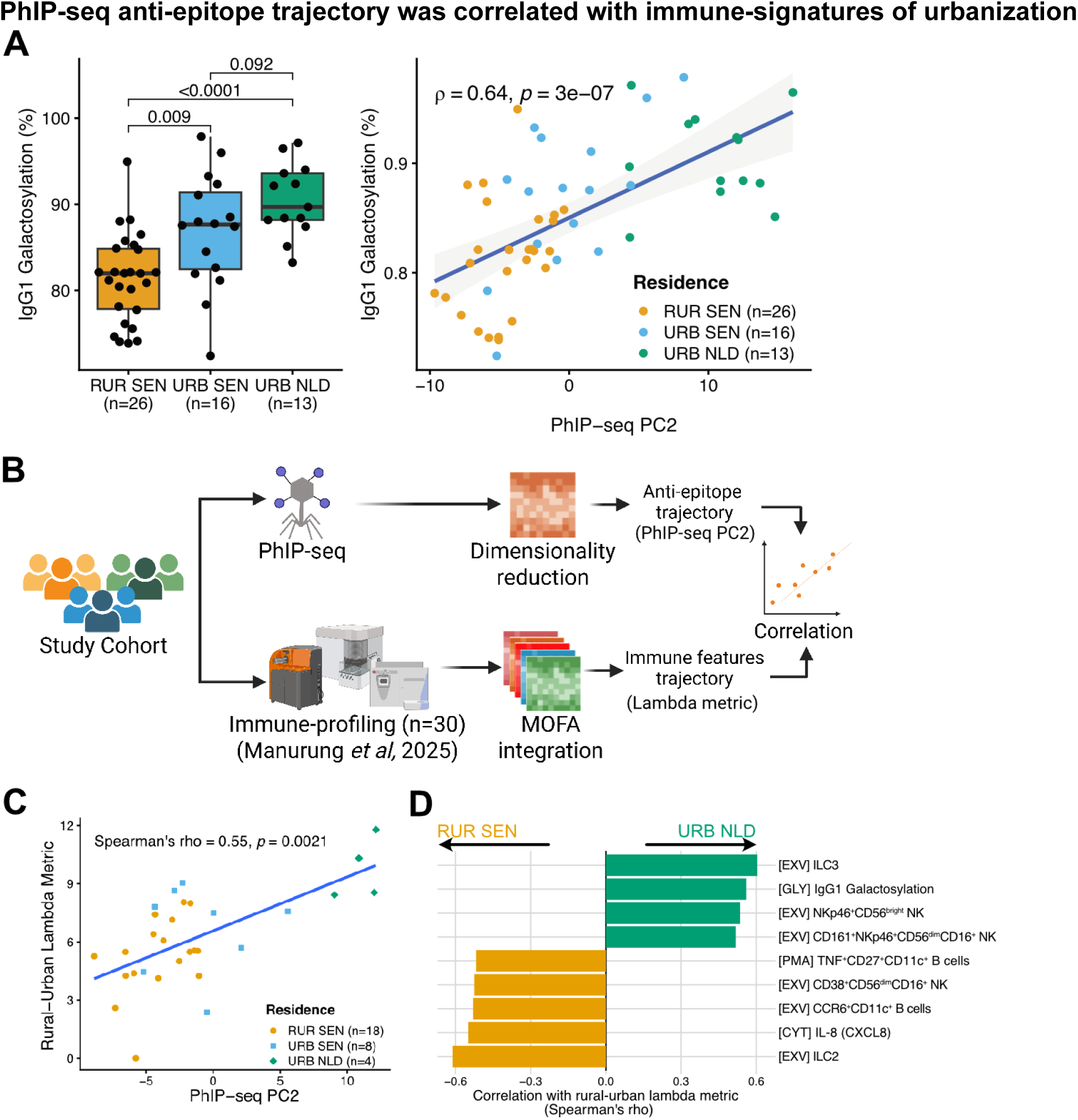
Associations between PhIP-seq anti-epitope trajectory and immunological parameters. A, Boxplots showing IgG1 galactosylation levels across the residence area (left); pairwise t-test p-values are shown. Scatter plot showing correlation between IgG1 galactosylation levels and PhIP-seq PC2 (right); Spearman’s correlation coefficient (ρ) and p-value are shown. **B**, Analysis scheme for integrating anti-epitope trajectory obtained from PhIP-seq (PhIP-seq PC2) with the previously published immune features trajectory obtained from immune profiling datasets (rural-urban lambda metric). Created using Biorender.com. **C**, Correlation between PhIP-seq PC2 scores and rural-urban lambda metric. Spearman correlation coefficient (rho) and *p-*value are shown with a fitted regression line. **D**, Immune features significantly correlated with PhIP-seq PC2 scores (FDR < 0.1), sorted by magnitude and direction of correlation coefficient.

To identify immune features driving this association, we correlated PhIP-seq PC2 scores with all immune parameters from our published dataset. Nine immune features exhibited significant associations (at FDR<0.1; Supplementary Fig. 6B), revealing distinct immunological signatures linked to exposure patterns. Rural-associated antibody profiles correlated with higher frequencies of ILC2s, CD11c^+^ B cells, and CD56^dim^CD16^+^ NK cells, as well as elevated serum IL-8 levels (Fig. 3D, Supplementary Fig. 6C). Conversely, urban-associated profiles correlated with increased ILC3 and NKp46^+^ NK cell frequencies alongside higher IgG1 Fc galactosylation. These findings demonstrate that serum antibody repertoires serve as robust proxies for microbial exposure histories and their downstream immunological consequences.

## Discussion

Using PhIP-seq, we profiled antibody repertoires of individuals across the rural-urban gradient and revealed distinct antigen exposure histories that correlate with environmental and lifestyle factors. Repertoires detected among rural Senegalese individuals were enriched for antibodies against pathogens associated with poor sanitation and endemic exposure, including *Shigella* species, human herpesviruses, and *Plasmodium falciparum.* Conversely common respiratory pathogens including Rhinovirus B, Epstein-Barr virus, and *Haemophilus influenzae*. These patterns align with known epidemiological differences in infectious disease burden between LMICs and HICs.

Interestingly, antibodies against human gammaherpesvirus 8–also known as Kaposi’s sarcoma herpesvirus (KSHV)–were detected among rural Senegalese participants but not their urban counterparts. KSHV is known to be highly prevalent in Sub-Saharan Africa and its higher prevalence in rural regions has been described previously (24). Studies have linked higher KSHV prevalence to factors typically associated with rural living, including denser household, which facilitate viral transmission (25), and *Schistosoma* infections, which induced viral reactivation (26)—factors also present in rural Senegal.

It is interesting to note that antibodies against certain epitopes are enriched in particular populations and that antibodies against epitopes from the same organism can display opposite enrichments. Enrichment of antibodies against particular epitopes has been reported in the context of disease, with antibodies against Epstein-Barr virus lytic phase proteins significantly enriched in individuals with primary sclerosing cholangitis compared to control (27) and antibodies against bacterial flagellin epitopes enriched in individuals with Crohn’s disease (12). For this paper, the antibody response *S. aureus* stands out, simultaneously detected in most individuals for some epitopes and for others enriched towards both ends of the rural-urban axis. The implications of these results, especially for vaccine development, is worth investigating further.

The differences in immune profile across the rural-urban axis is hypothesised to be driven, at least in part, by pathogen exposure in the environment (7, 23), as suggested by differences in infectious disease burden between LMICs and HICs (28, 29). Here, we demonstrated the convergence between the independently constructed trajectories of antibody repertoire profiles and immune remodeling along the rural-urban gradients. This analysis further revealed specific immune features associated with rural-urban exposure histories. For example, ILC2s, which was associated with rural repertoire profiles, were known to be expanded following helminth infections (17). Conversely, urban repertoire profiles were associated with higher frequencies of NK cells expressing NKp46, critical NK cell marker for controlling viruses such as influenza (30). Future studies with larger sample size will provide greater statistical power to link specific microbial epitopes to immune parameters to further our understanding of mechanistic link between exposures and specific factors driving it.(28, 29). Together these findings highlight the need to utilise this technology to further investigate the link between pathogen exposures and immune adaptations.

This work can be extended upon in multiple ways, the first being to increase sample size to numbers similar to other papers (12, 22), allowing greater statistical rigour and overcoming interpretation issues confounded by individual library size, such as exposure diversity and read count levels. The second expansion would be in the pathogen epitopes investigated. Most of the epitopes are from microbiome organisms, reflecting the background of PhIP-seq in studying inflammatory bowel diseases (12, 21). When assessing differences across the rural-urban axis it would be beneficial to include epitopes from pathogens common to LMICs, but not HICs, such as helminths (29). Nevertheless, the fact that most of our findings align with epidemiological observations, such as higher detection of KSHV in rural Senegal, highlight the strength of PhIP-seq as a tool for a broad, systematic survey of microbial exposures and might inform regional disease screening policies.

In conclusion, PhIP-seq robustly captures differential microbial exposure patterns across rural-urban gradients, providing insights into how environmental exposures shape population-specific immune signatures. This approach establishes a framework for understanding immune adaptation in diverse global populations and environments.

## Materials and methods

### Study design

Participants were recruited between November 2018 and August 2019 from three sites: rural Senegal (Pakh and Richard Toll; n=72), urban Senegal (Dakar; n=46), and the Netherlands (Leiden; n=28). Recruitment was coordinated through local healthcare centers in Senegal and Leiden University Medical Center. Written informed consent was obtained from all participants. The study was approved by the ethics committees of Cheikh Anta Diop University of Dakar (0339/2018/CER/UCAD) and Leiden University Medical Center (NL66287.058.18, NL- OMON48968).

Rural Senegalese participants originated from agricultural villages in the Senegal River basin, where schistosomiasis has historically been highly endemic. Urban Senegalese participants were recruited from Dakar, a non-endemic area with improved sanitation and consistent access to clean water. Retrospective surveys in Dakar reported intestinal parasite prevalence of ∼19– 26%, although estimates may be biased by sampling from symptomatic patients. Rural and urban classifications were confirmed using questionnaire data on household size, animal contact, housing conditions, and amenities. The Senegalese population is ethnically diverse (predominantly Wolof, Pular, and Serer), though cultural admixture is common; most participants were Wolof.

Inclusion criteria were: age 18–40 years, residence in the study area ≥10 years, and absence of chronic illnesses (e.g. diabetes, hypertension). Participants underwent clinical examination and completed standardized questionnaires covering demographic, socioeconomic, lifestyle, and medical history information. Exclusion criteria were evidence of infection with *Plasmodium spp.* (thick smear, rapid diagnostic test), *Schistosoma spp.* (Kato-Katz and urine filtration, 12 µm filters), or intestinal helminths (*Ascaris lumbricoides*, *Trichuris trichiura*, hookworm) as determined by stool microscopy.

To ensure comparable sample quality across sites, technical staff received centralized training at Leiden University Medical Center and adhered to standardized operating procedures (SOPs) for sample collection and processing. Venous blood was drawn into heparin, EDTA, and dry tubes for cell isolation, full blood count, and serology testing.

### Description of immune-profiling datasets

Five datasets from our previously published study (7) were re-analyzed in this manuscript: peripheral blood immune cell phenotyping (*ex vivo*; EXV), cytokine production following 6-hour stimulation with PMA/ionomycin or monophosphoryl lipid-A (MPL), IgG Fc glycan profiling (GLY), and serum cytokine levels (CYT). Detailed methodological description for the acquisition, pre-processing, and analysis for each dataset as well as the integrative analysis of these datasets can be found elsewhere (7).

Briefly, the EXV, PMA, and MPL datasets were obtained from mass cytometry analysis of peripheral blood mononuclear cells (PBMCs). Cryopreserved PBMCs (3×10⁶ cells) were cultured with either PMA/ionomycin or MPL for 6 hours (Brefeldin-A added for the final 4 hours), then stained using two standardized antibody panels for surface markers, intracellular cytokines, and nuclear antigens, with dead cell discrimination via intercalator staining. Cells were acquired on a Helios mass cytometer with automated tuning, normalization using calibration beads, and quality control through FlowJo gating and specialized R packages. Batch effects in stimulated samples were corrected using CytoNorm, followed by two-step unsupervised clustering via FlowSOM and FastPG to identify immune cell clusters.

IgG Fc-glycopeptides (GLY) were profiled by affinity purification on Protein G beads, trypsin digestion, and LC–MS analysis. LacyTools was used for glycopeptide quantification, yielding subclass-specific Fc glycosylation profiles for IgG1 (24 glycopeptides) and IgG2 (14 glycopeptides); IgG3/IgG4 peptides were excluded due to allotype overlap. From these data, glycosylation traits—including fucosylation, galactosylation, sialylation, bisection, and the sialic acid/galactose ratio—were derived using standard formulas based on relative glycopeptide abundances.

Plasma cytokines (CYT) were quantified using the R&D Systems Luminex Discovery Assay Human Premixed Multi-Analyte Kit (CCL3, CCL8, CCL24, complement component C5a, CXCL10, HGF, IL-3, IL-8, IL-18, vascular endothelial growth factor, CCL2, CCL7, CCL20, CXCL9, CXCL13, IFN-γ, IL-6, IL-10, and S100A8) with reduced reagent volumes. Analytes undetectable in >40% of samples were excluded, and values below the lower limit of detection (LLOD) were imputed as half the LLOD.

Finally, these immune-profiling datasets were integrated using MOFA2 R package–an unsupervised method that learns latent factors capturing variations across datasets. Latent factors were further reduced into two dimensions using multidimensional scaling and then used as input for trajectory analysis using principal curve (princurve R package), which have previously revealed a trajectory of immune variation across the rural-urban gradient.

### Phage immunoprecipitation sequencing (PhIP-Seq)

PhIP-Seq assays were conducted following the protocol described by Vogl et al. (2021), using the same core parameters with a few minor modifications: a 4,000-fold coverage of phages per variant and 2 µg of antibodies per reaction. The microbiota library, consisting of 244,000 variants, was mixed with two additional libraries: a 100,000-variant pool containing peptides derived from allergen and environmental compound databases (as described by Leviatan et al.) (31), and a 13,192-variant pool primarily composed of Coronavirus antigens (developed by Klompus et al.) (32). Bead washing steps were automated using a Tecan Fluent 1080 liquid-handling robot. PCR amplifications were performed using primers specified in Vogl et al. (2021) (21), except for PCR2, where 10-nucleotide unique dual index primers (purchased from IDT) were used.

### PhIP-Seq data processing

Samples were sequenced with Illumina Novaseq X. Raw reads bam files were then converted to fastq format using bamtofastq (bedtools v2.31.0) and subsequently analyzed through a previously established workflow. Reads were mapped to the three antigen libraries mentioned above [refs], and raw counts for each peptide were obtained. To assess the enrichment of antibody bound peptides, peptide counts were normalized by comparing them to the oligopeptide counts in the initial library as previously done. The phage library was sequenced before immunoprecipitation and peptide counts were modeled as a generalized Poisson distribution to form the null model. For each sample, peptide enrichment significance (p-values) was computed while controlling for Family Wise Error Rate (FWER) with the Bonferroni method.

### Annotation of epitopes

The agilent library peptides were run against the Uniref90 (33) database to annotate the protein and organism of origin. Some of the results from the Uniref90 database did not have a specific tax, showing up as “Bacteria” or “root”. The amino acid sequences from the non-specific agilent and the entire twist peptide libraries were analysed with blastP against non-redundant protein sequences and the result XML files retrieved. For each epitope, the scientific names of the top 15 results (in terms of E-value) were extracted. The organism origin of the epitope was assigned to be the most common scientific name in the top 15 list. Ties were broken randomly. Manual standardization of the organism names was also carried out, for example removing strain information from the scientific name and ensuring continuity between the names derived from the Uniref90 database and blast. For example, cytomegalovirus is named “Human betaherpesvirus 5” in blast results and “Human cytomegalovirus” in the Uniref90 results. This was standardised to “Human cytomegalovirus”.

### Statistical analysis

All statistical analyses were performed in R (v4.4.1). All data points represent one biological replicate, and technical replicates were not performed. No sample points were omitted from analysis. All testing was two sided.

The PhIP-seq dataset was treated as a binary matrix showing presence/absence of antibody against a particular epitope. Individuals who had a significant enrichment for a particular epitope were treated as though an antibody against that epitope was detected in the individual.

Principal component analysis was performed using the R stats package on a subset of the binary PhIP-seq dataset where epitopes were removed that were expressed in less than 25% of the individuals across all three residence groups. Significant pairwise differences between residence and sex groups in the PC space were assessed using PERMANOVA, as implemented in the adonis2 function of the Vegan R package.

To assess whether the frequency of epitopes showed a linear trend across the residence groups a Cochran-Armitage test was performed across all the individuals and epitopes, with p-values adjusted using Benjamini-Hochberg procedure to account for multiple testing. For testing linear trend across the continuous data, orthogonal polynomial contrast was utilised where features which were significant for linear (but not quadratic) trend were treated as significant.

## Acknowledgements

We thank the laboratory members of the M.Y. group for their critical feedback. We would like to acknowledge all staff members in Senegal (field workers, medical staff of the health Centre of Richard Toll, staff of the Immunology Laboratory at Cheikh Anta Diop University, Senegal) and the Netherlands who helped to make this study possible. Last, we would like to thank all volunteers who participated in this study, without whom the study could not have been performed.

## Author contributions

M.Y., M.M., and M.D.M. were responsible for the study design. R.F.L. and M.D.M. were responsible for the statistical analysis and data interpretation and prepared the first draft. M.Y., M.M., and M.D.M. acquired funding. G.I. and T.V. generated the data and optimized the PhIP-seq experimental protocols. All authors reviewed the manuscript.

## Data and materials availability

Data are available upon reasonable request to the authors.

## Supplementary information

**Supplementary Figure 1.**
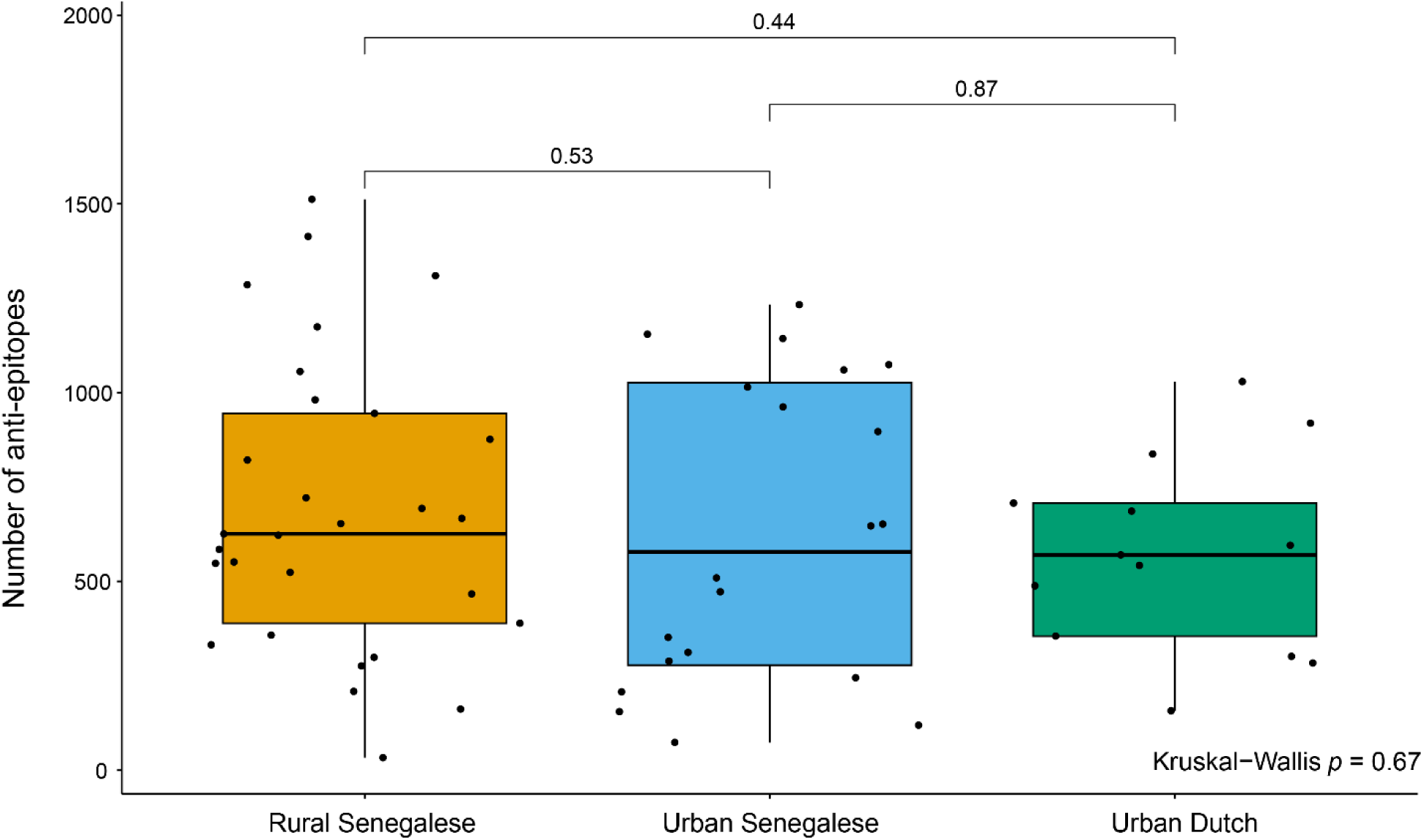
Boxplot showing the number of significant anti-epitopes identified for each individual, grouped by the residence of individual. Kruskal Wallis and pairwise Wilcoxon Rank Sum p values are indicated.

**Supplementary Figure 2.**
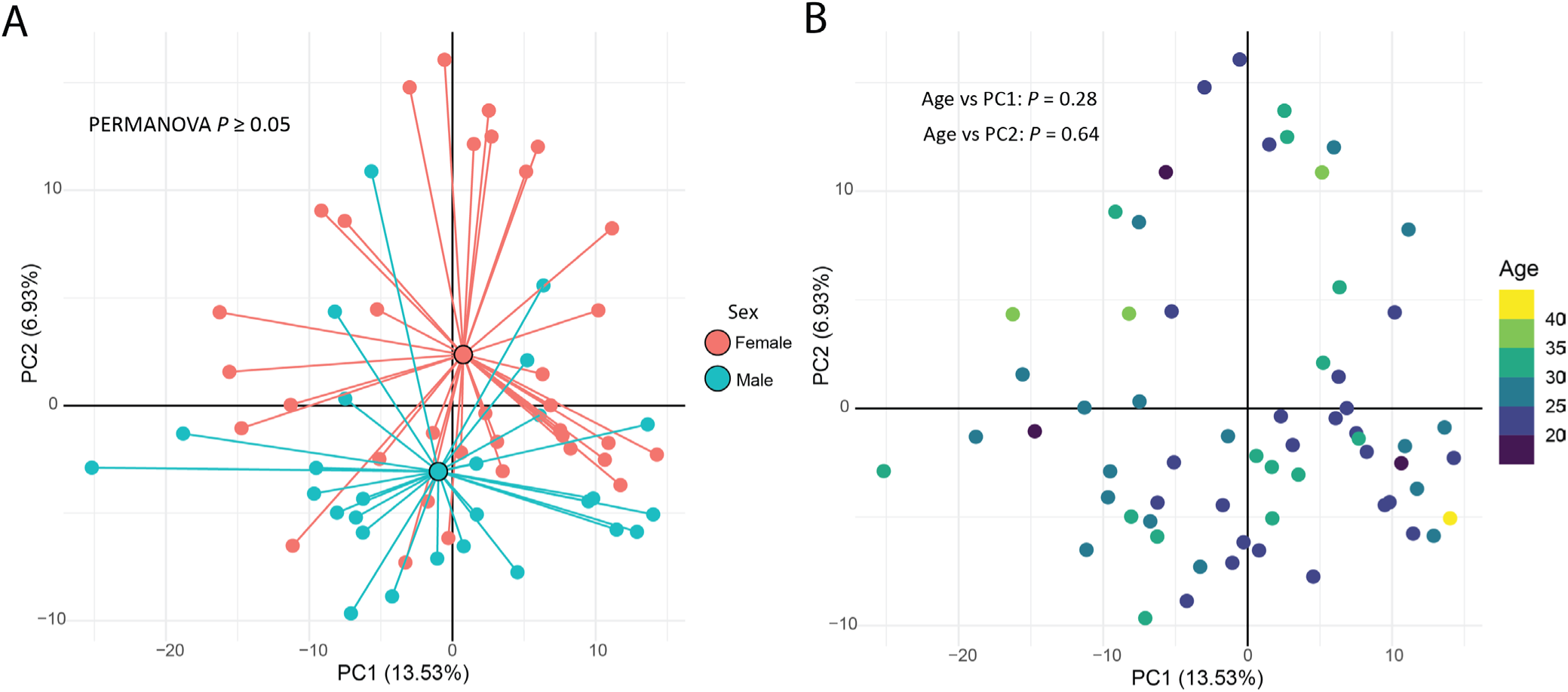
PCA plot generated from the filtered PhIP-seq dataset, where individuals are coloured by sex (**A**) and age (**B**). The large circles represent the centroid of the different groups in the PC1 and PC2 space. *P* values denote group centroid comparisons with the PERMANOVA test (left) or Pearson correlation test (right).

**Supplementary Figure 3.**
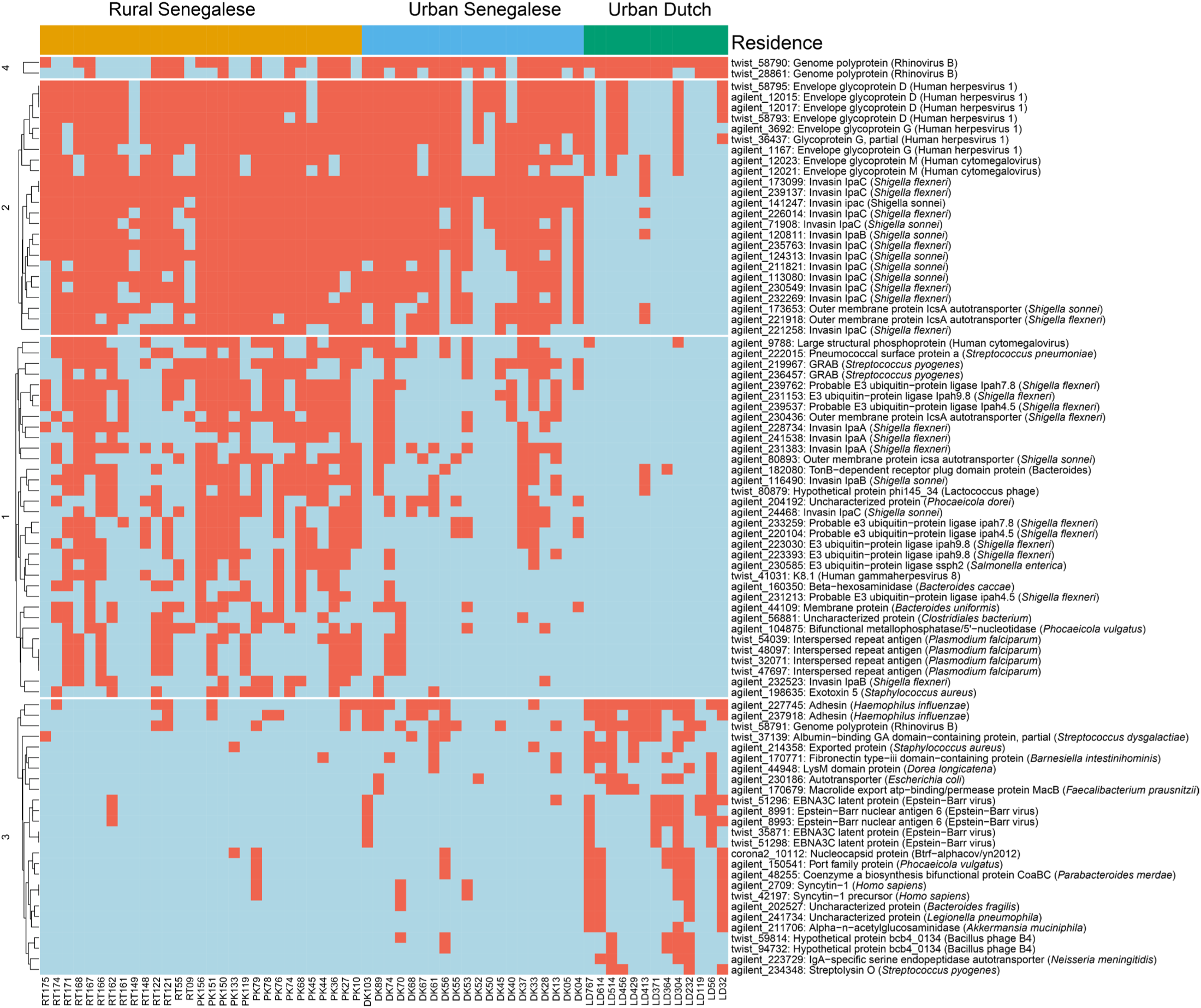
Clustered heatmap showing whether antibody against a particular epitope was detected (red) or undetected (blue) in an individual. The heatmap is sliced column wise by the residence of the individual, and the rows sliced by k-means clustering on the epitopes. Epitope IDs, protein and organism of origin are annotated on the rows and are coloured according to their type, as in Figure 2A & B.

**Supplementary Figure 4.**
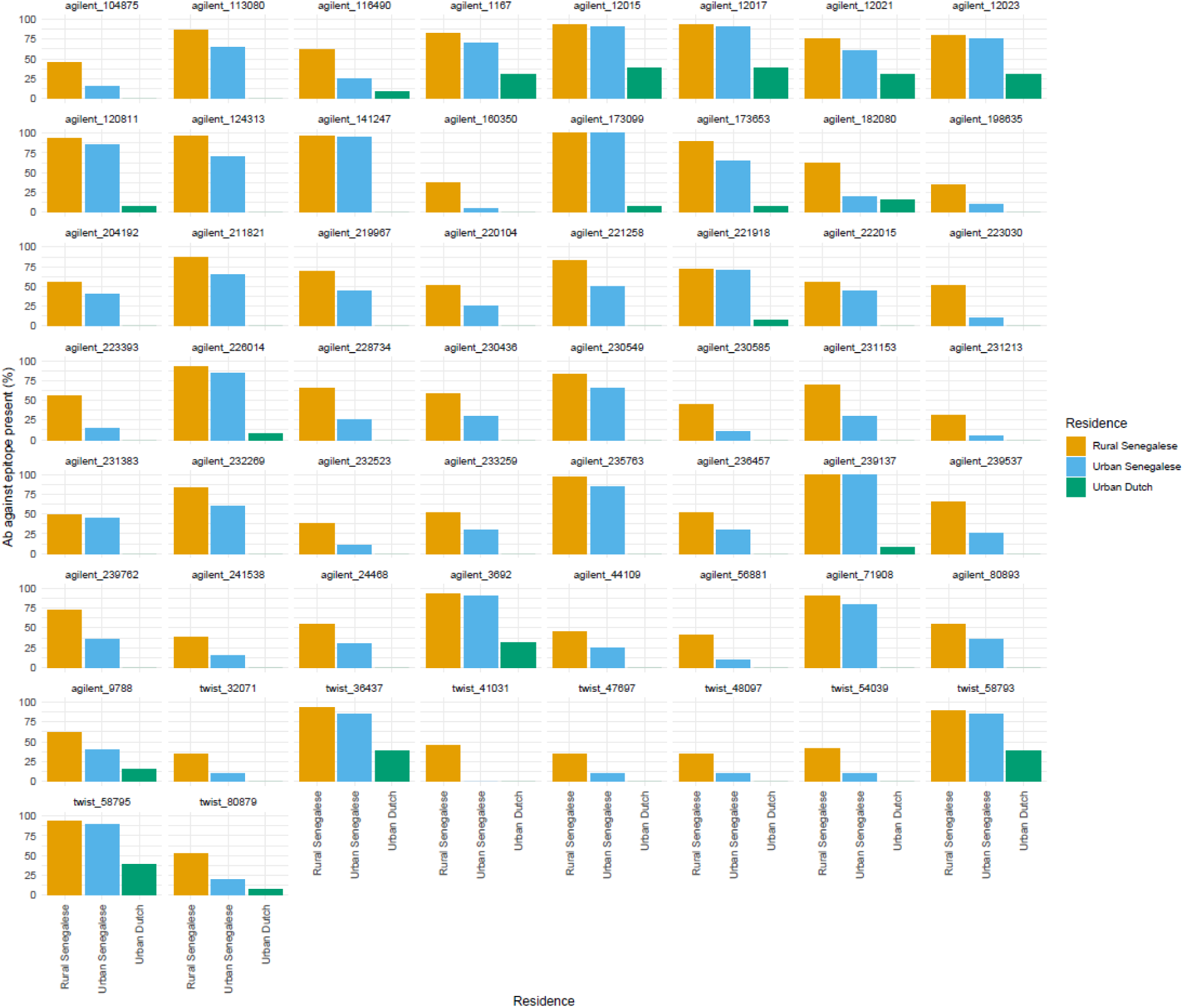
Barplots of all the anti-epitopes which had a significant linear trend towards the rural Senegalese side of the rural-urban axis. Barplots show the percentage of individuals per group who have antibodies against that particular epitope

**Supplementary Figure 5.**
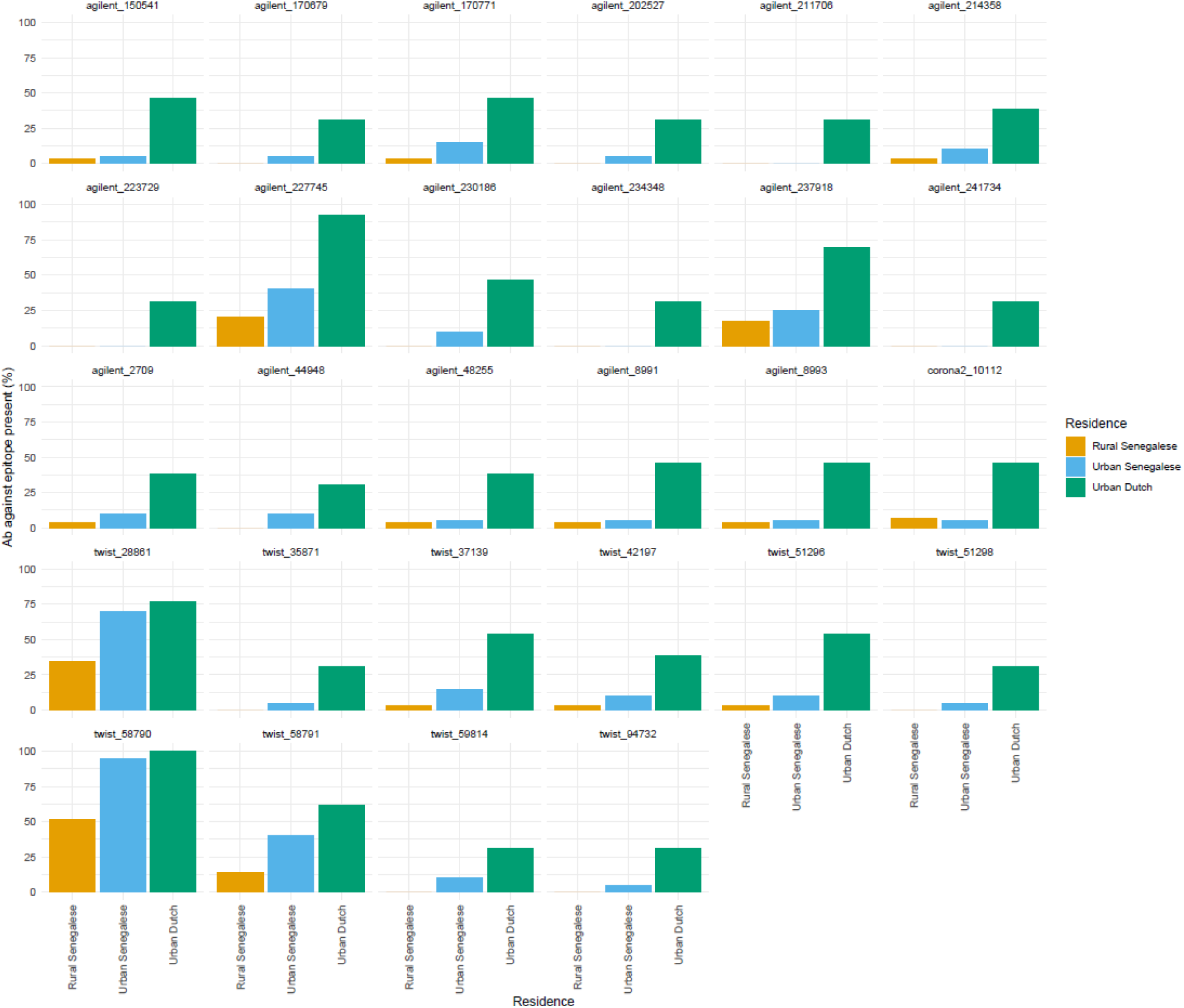
Barplots of all the anti-epitopes which had a significant linear trend towards the urban Dutch side of the rural-urban axis. Barplots show the percentage of individuals per group who have antibodies against that particular epitope

**Supplementary Figure 6.**
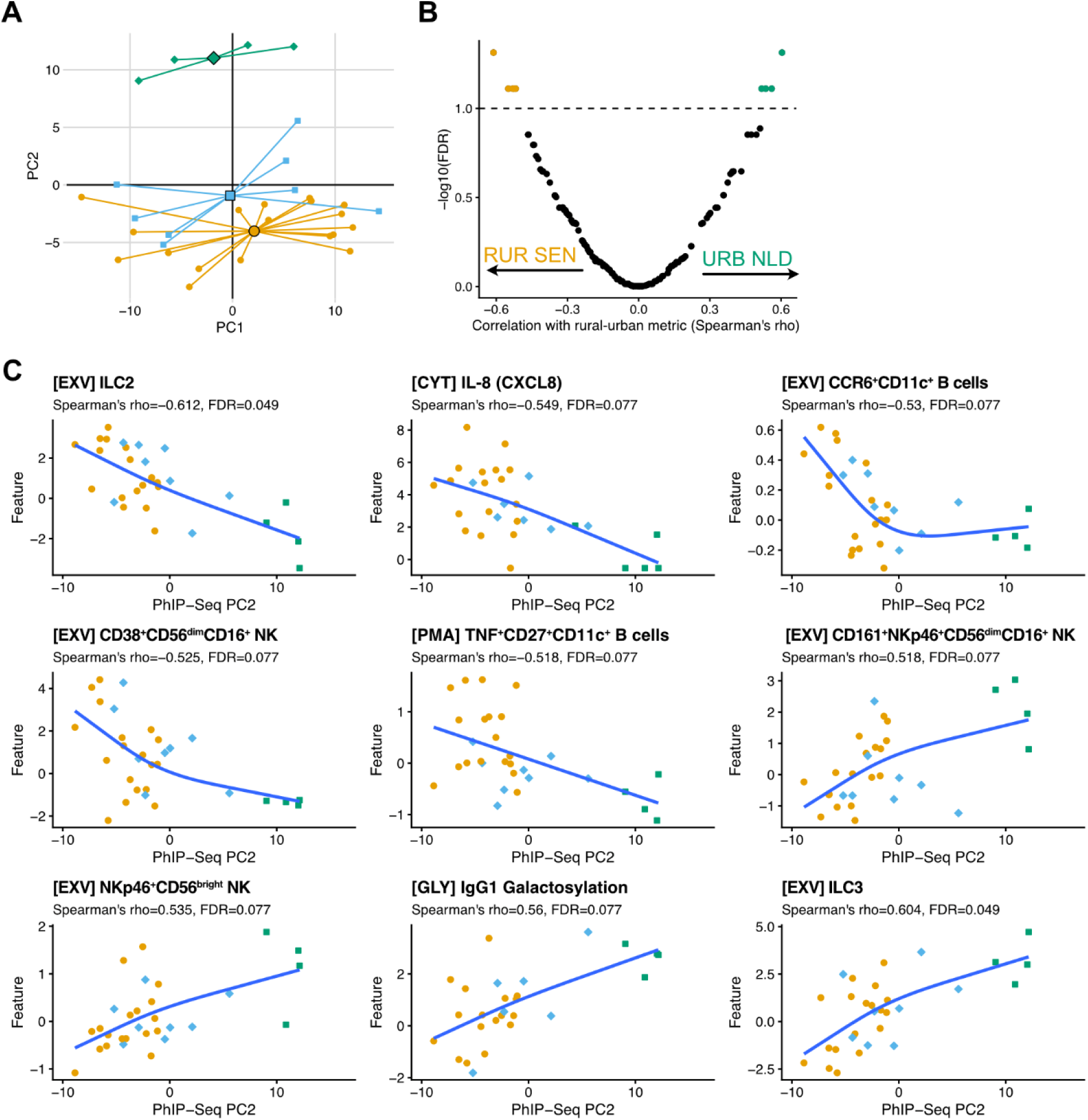
**A**, PCA plot of PhIP-seq dataset, subsetted for individuals with corresponding high-dimensional immune-profiling datasets. **B**, Volcano plot of correlations between PhIP-seq PC2 trajectory with immune parameters. **C**, Scatter plot showing immune parameters with statistically significant correlation (FDR<0.1) with PhIP-seq PC2 trajectory.

Supplementary File 1.

File containing all epitopes in which antibodies were detected in at least one individual. Information included is whether the organism name was derived from blastP or not, the number of individuals in which antibody was detected, the amino acid sequence and the epitope ID.

Supplementary File 2.

File containing all the epitopes which were found to have a significant linear trend across the rural-urban axis. Statistics from the Cochran-Armitage test results can be found, as well as the epitope ID and the protein and organism from which the epitope is from.

